# Examination of Australian *Streptococcus suis* Isolates From Clinically Affected Pigs in a Global Context and the Genomic Characterisation of ST1 as a Predictor of Virulence

**DOI:** 10.1101/375543

**Authors:** Mark A O’Dea, Tanya Laird, Rebecca Abraham, David Jordan, Kittitat Lugsomya, Laura Fitt, Marcello Gottschalk, Alec Truswell, Sam Abraham

**Affiliations:** Antimicrobial Resistance and Infectious Disease Laboratory, School of Veterinary and Life Sciences, Murdoch University, Perth, Western Australia; Wollongbar Primary Industries Institute, NSW Department of Primary Industries, NSW, Australia; Department of Veterinary Microbiology, Faculty of Veterinary Science, Chulalongkorn University, Bangkok, Thailand; ACE Laboratory Services, Bendigo, Victoria, Australia; Laboratory of Research on *Streptococcus suis*, Faculty of Veterinary Medicine, University of Montreal, Saint-Hyacinthe, QC, Canada

**Author notes:** address correspondence to Mark O’Dea; Sam Abraham.

## Abstract

*Streptococcus suis* is a major zoonotic pathogen that causes severe disease in both humans and pigs. In this study, we investigated *S. suis* from 148 cases of clinical disease in pigs from 46 pig herds over a period of seven years. These isolates underwent whole genome sequencing, genome analysis and antimicrobial susceptibility testing. Genome sequence data of Australian isolates was compared at the core genome level to clinical isolates from overseas. Results demonstrated eight predominant multi-locus sequence types and two major *cps* gene types (cps2 and 3). At the core genome level Australian isolates clustered predominantly within one large clade consisting of isolates from the UK, Canada and North America. In particular, serotype 2 MLST25 strains were very closely associated with Canadian and North American strains. A very small proportion of Australian swine isolates (5%) were phylogenetically associated with south-east Asian and UK isolates, many of which were classified as causing systemic disease, and derived from cases of human and swine disease. In addition, we show that ST1 clones carry a constellation of putative virulence genes not present in other Australian STs, and that this is mirrored in overseas ST1 clones. Based on this dataset we provide a comprehensive outline of the current *S. suis* clones associated with disease in Australian pigs and their global context, and discuss the implications this has on antimicrobial therapy, potential vaccine candidates and public health.

**Importance:** In this study, we examine in detail, the genomic characteristics of 148 *Streptococcus suis* isolates from clinically diseased Australian pigs. We report the antimicrobial susceptibility profiles, virulence gene analysis and relationship to isolates from other regions of the world. We also demonstrate that ST1 clones, regardless of serotype, carry a large array of putative virulence genes while maintaining a small total gene content. This compilation of data has major ramifications for vaccine development, and refines the understanding of the distribution of various strains of this potentially-fatal zoonotic agent in the global pig industry

## Introduction

*Streptococcus suis* is an agent of serious concern for the global swine industry, and an emerging zoonotic agent causing meningitis, sepsis, arthritis, endocarditis and endopthalmitis in humans. Human disease is particularly prevalent in the western Pacific and south-east Asian regions, where risk is predominantly attributable to poor sanitation at slaughter and ingestion of undercooked pork, followed by Europe and then significantly less in the Americas (1). Zoonotic infection with *S. suis* has occurred in Australia, although there are few reports in the literature. The most recently published cases occurred in 2007 and 2008 in an abattoir worker from Victoria and two piggery workers from New South Wales respectively (2, 3) with each case attributed to serotype 2, which, along with serotype 14, is one of two globally-dominant serotypes detected in human cases.

In swine, *S. suis* is predominantly carried on the tonsils, but also within nasal cavities, genital and gastrointestinal tracts of many clinically healthy pigs, a feature which complicates management strategies in herds with disease outbreaks. Despite this carriage in healthy pigs, *S. suis* is a cause of a wide range of clinical syndromes ranging from sudden death in the peracute manifestation of disease, to meningitis, septicaemia, endocarditis, arthritis and pneumonia. The clinical manifestations generally occur in the post-weaning period, although can occur less frequently in suckers and in adult pigs (4). Isolates obtained from areas indicative of invasive disease such as joints, brain tissue, heart and abdomen may be referred to as systemic, or causative of systemic infection, in contrast to isolates obtained from lung tissue.

Treatment for affected animals is usually reliant on administration of β-lactams such as penicillin and amoxicillin, and, where permitted, farms may prophylactically treat all pigs at the peri-weaning stage. Another management option employed by some producers is vaccination, usually in the form of bacterins produced as autogenous vaccines from on-farm isolates (5). Due to the highly variable antigenicity of the capsular polysaccharides (CPS), *S. suis* is currently classified into 29 serotypes (six previously classified serotypes have been reassigned to different bacterial species) (6), and it is considered that protection (if any) is only provided by homologous vaccine serotypes. In addition to this, virulence factors and yet to be determined genetic factors may play a role in poor vaccine efficacy (7), although virulence factors are not necessarily protective antigens and the current uncertainty surrounding the role of virulence factors makes this difficult to assess.

A recent review on the worldwide distribution and typing of *S. suis* by Goyette-Desjardins et al. (2014) provided a comprehensive analysis of the predominant serotypes and MLSTs in swine production systems across the US, South America, Europe and South-East Asia (8). Of note in this review was the lack of information on Australian *S. suis* types, with no published surveillance since 1994 (9), a gap of 20 years. Following this a study published in 2015 on 45 Australian *S. suis* isolates (encompassing 3 human isolates from 2006-2008, and 42 swine isolates from 1981-2011) described 4 MLST’s (1, 25, 369 and 28) (10).

Given the paucity of data available on *S. suis* associated with disease in Australian pigs, the aim of this study was to analyse and characterise a significant number of *S. suis* isolates from diseased pigs obtained across multiple production sites and spanning a seven year period. In addition to serotype and MLST, we report on virulence factors and antimicrobial susceptibility, and compare the core genome to international isolates to assess evolution in a global context.

## Results

### Samples

A total of 148 swabs from archived clinical isolates were transferred from ACE Laboratory Services (a major swine-industry referral lab) to the Murdoch University Antimicrobial Resistance and Infectious Diseases Laboratory for detailed molecular analysis. The majority of isolates were acquired from the lungs of diseased pigs and were classified as potentially virulent, followed by isolates from heart, brain, abdomen and joints which were classified as virulent isolates, and miscellaneous sites including upper respiratory tract, abscesses and lymphoid tissue (Table 1). Isolates spanned a seven year period with 2, 1, 29, 27, 9, 49 and 29 isolates from years 2010, 2011, 2013, 2014, 2015, 2016 and 2017 respectively, and two isolates for which the year was unknown. Isolates were originally obtained from at least 10 of Australia’s major pig production enterprises (17 samples were of unknown origin) and encompassed 46 known individual farms (7 isolates had unavailable farm data). Of the available farm data, 39 farms accounted for between one and five isolates each, five farms accounted for between six and eight isolates each and one farm accounted for 33 isolates.

**Table 1.** Number of isolates and site of origin used in the study

| Isolation Site | Number of Isolates |
| --- | --- |
| Abdomen | 2 |
| Brain | 12 |
| Heart | 12 |
| Joint | 6 |
| Lung | 93 |
| Lymph node | 1 |
| Neck abscess | 3 |
| Upper respiratory tract | 3 |
| Unknown | 16 |

### Genomic and phylogenetic analysis

Genomic analysis of the 148 isolates revealed that 110 isolates belonged to 11 previously identified MLSTs with 38 isolates belonging to one of 26 newly identified MLSTs. The most prominent MLSTs were ST 27 (27/148), ST 25 (26/148), ST 28 (11/148), ST 483 (11/148) and ST 1 (10/148). The new MLSTs assigned as a result of this study were ST 1031 – ST 1056 inclusive.

Analysis of the capsular genes resulted in two major serotypes being detected; serotype 2 (39/148) and serotype 3 (37/148). The other main serotypes were 1/2 (11/148), 16 (8/148), 19 (8/148), 8 (7/148) and 4 (7/148). Analysis of serotypes and MLST combined showed a high proportion of isolates as serotype 2 ST25 (17.6%), serotype 2 ST28 (6.1%) and serotype 3 ST27 (18.2%). Of serotypes where more than one isolate was present, serotype 15 (n=2), 16 (n=8), 21 (n=4) and 31 (n=5) all had previously unidentified STs.

The proportion of isolates carrying the main virulence factors *mrp, epf* or *sly* across all isolates was 7.4% (cps 1/2, 2, 14 and 19), 6.8% (cps 1/2, 2 and 14) and 33.1% (caps1/2, 2, 3, 4, 5, 7, 8, 11, 12, 14, 15, 16, 18, 19, 21, and 23) respectively. Only a single isolate carried both *mrp* and *sly,* while 6.8% of isolates carried both *mrp* and *epf,* and 6.8% carried *mrp, epf* and *sly.* The isolates carrying all three genes belonged to serotypes 1/2 (n=7), 2 (n=2) and 14 and all belonged to MLST 1, indicating a significant association between ST 1 and the carriage of all three genes (Fisher’s exact test P-value <0.0001). Although predominantly isolated from lung, two of these isolates originated from brain and two from joint fluid and all four of these isolates were serotype 1/2. These serotype 1/2 ST1 isolates also carried more virulence genes with an average of 46 compared to all other isolates carrying 36 virulence genes and were clustered in virulence gene group 2 (Figure 1).

**Figure 1:**
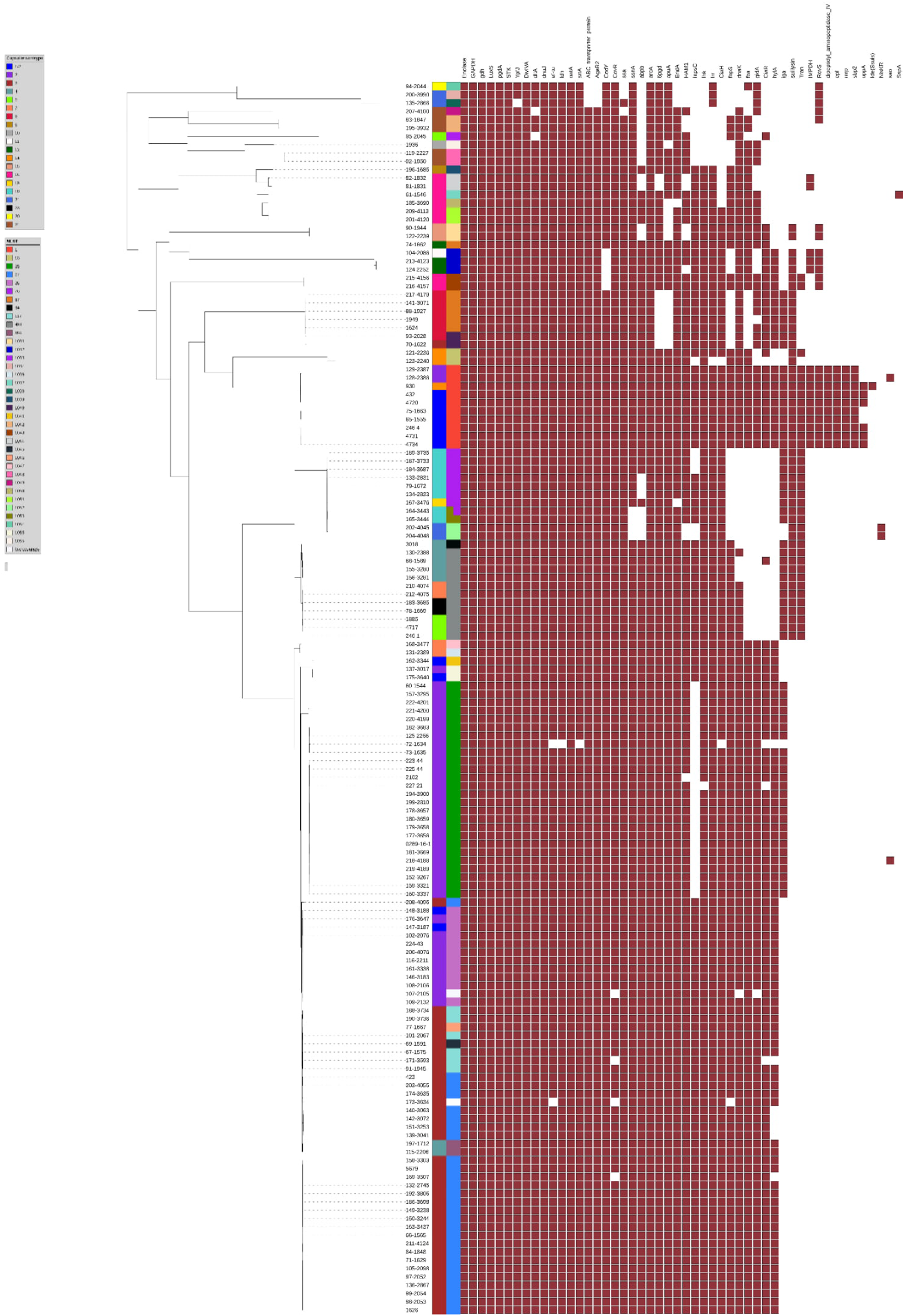
Presence of putative virulence factors in all 148 isolates from this study. Blocks indicate presence of individual genes. Genes are identified by columns and isolate number, MLST and serotype are identified in rows.

The main putative virulence genes present across all isolates included *enolase* (100%), *GAPDH* (100%), *gdh* (100%), *luxS* (100%), *pgdA* (100%), *stk* (100%), *divIVA* (99.3%), *dltA* (99.3%), *dnaJ* (99.3%), *ef-tu* (99.3%), *ldh* (99.3%), *oatA* (99.3%), *srtA* (99.3%) and *ypfJ* (99.3%) and this conserved group was the only common feature found in isolates associated with systemic disease (brain, joint, heart, abdomen isolates). Detailed examination of the virulence gene profiles demonstrates that the Australian isolates can be grouped according to four main blocks of virulence genes (Figure 1). Aside from the core virulence genes present in most isolates, group 1 can be broken into subgroups based on small clusters of genes, however no discernible pattern relating to potential virulence is present. Group 2 contains a conserved block of 45 virulence genes and is the only group to contain the genes *dppIV, epf, mrp, sbp2, oppA* and *lde*. This group consists predominantly of serotype 1/2 and serotype 2 isolates, and all are ST1. Groups 3 and 4 are similar in gene composition with the exception that group 3 isolates are lacking a combination of *fba, gidA, ciaR* and *hylA,* and group 4 isolates have a combination of *iga, sly* and in some cases *fbps* and *dnaK.* However group 3 isolates have a combination of *iga, sly* and *Tran* which is not present in Group 4.

Analysis of the amino acid sequence of GDH across the Australian strains showed that all ST1 isolates had amino acid substitutions of A, S, K and K at positions 296, 299, 305 and 330 respectively, whereas non-ST1 isolates had the substitutions A,A,E,E, or T, A, E, E at the corresponding positions.

On average, the total gene content of systemic isolates from brain and joint, was 2057, significantly lower than that from respiratory isolates which carried 2109 (Mann-Whitney *U* test p-value = 0.025). When isolates from brain, joint, heart and abdomen were analysed, the average total gene content was 2095, which was not significantly different from respiratory isolates (p-value=0.682). When the total gene content of isolates was compared to the number of virulence genes a trend was observed based upon ST (Figure 2a). This was most apparent in the ST1 isolates which had a significantly higher number of virulence genes compared to the other ST’s, and the lowest number of total genes with an average total gene content of 1918 (p-value<0.00001). ST25 isolates also formed a tight cluster with a mean of 35.7 virulence genes and a relatively low total mean gene content of 2050. Despite having a mean virulence gene content of 36, ST28 isolates clustered with some of the highest total gene contents with all isolates having greater than 2100 genes and 9/11 having greater than 2200 genes. These findings were mirrored by an assessment of global *S. suis* virulence genes against total gene content (Figure 2b). Most striking is the consistent clustering of ST1 isolates with a mean virulence gene content of 45.8 and a mean gene content of 1936. The six non-ST1 isolates within this cluster consist of ST107 (n=1), ST105 (n=2) and ST144 (n=3) which are single locus variants (SLV) of ST1 at the *dpr, gki* and *cpn60* loci respectively. Additionally, for the published systemic isolates from Weinert et al. (2015), from which data was analysed and MLST able to be determined using our pipeline, 51/108 (47.2%) were ST1, with the next highest proportion being ST28 at 17/108. When SLVs were analysed, another seven isolates were found to be variants at the *gki* locus, such that when ST1 and ST1 SLV isolates were combined they accounted for 53.7% of systemic isolates.

**Figure 2.**
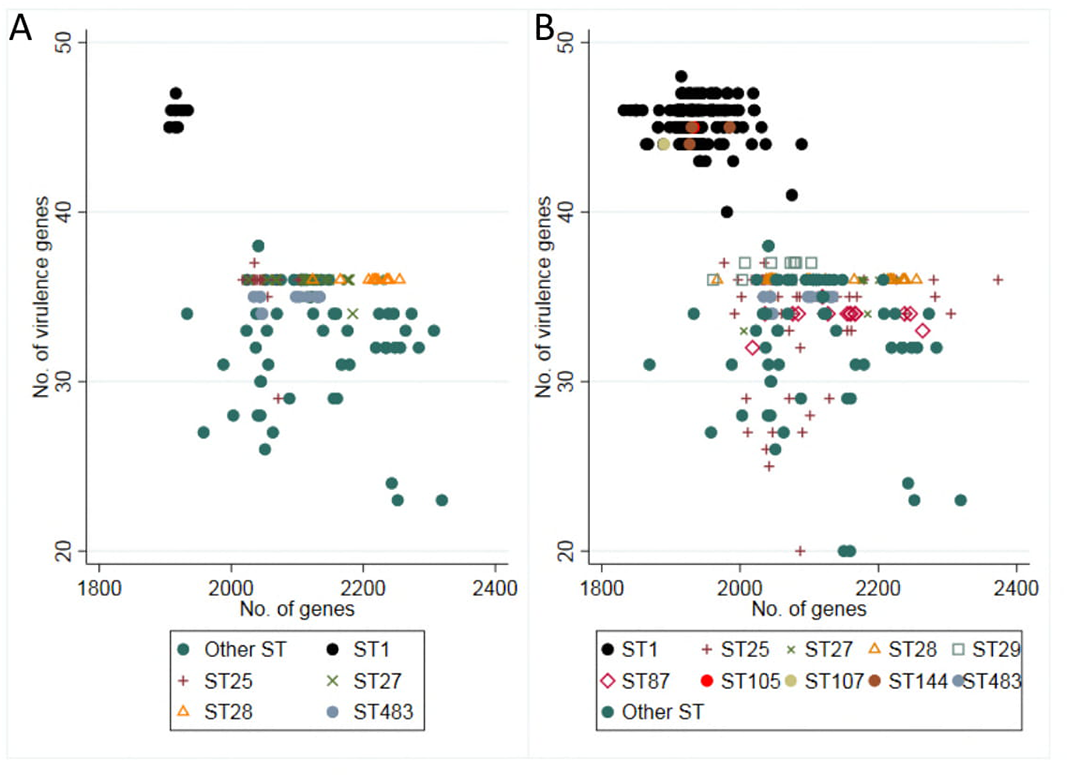
**a)** Total gene content of individual Australian isolates plotted against virulence factor content based on MLST. **b)** Total gene content of international isolates for which MLST could be determined, plotted against virulence factor content.

Phylogenetic comparison of the *S. suis* isolated from Australian pigs against a global collection of *S. suis* strains resulted in the identification of four major clades (Figure 3). The strains from this study were present in all four clades. Clade 1 was highly divergent from the other clades, and contained 20 Australian strains, with strain 74-1662 (serotype 12 ST87), an isolate from lung tissue, being the most divergent. The majority of the Australian strains (89/148) clustered in clade 2, which was comprised entirely of strains from the UK, North America and Canada. Only eight strains were present in clade 4, which was made up of Vietnamese and UK isolates. Australian serotype 2 ST25 isolates clustered with Canadian and North American serotype 2 ST25 isolates within clade 2. To explore this relationship, further analysis of the core genome was performed on these isolates (Figure 4) demonstrating that despite the similarity, the Australian isolates can be distinguished from North America and Canadian strains.

**Figure 3.**
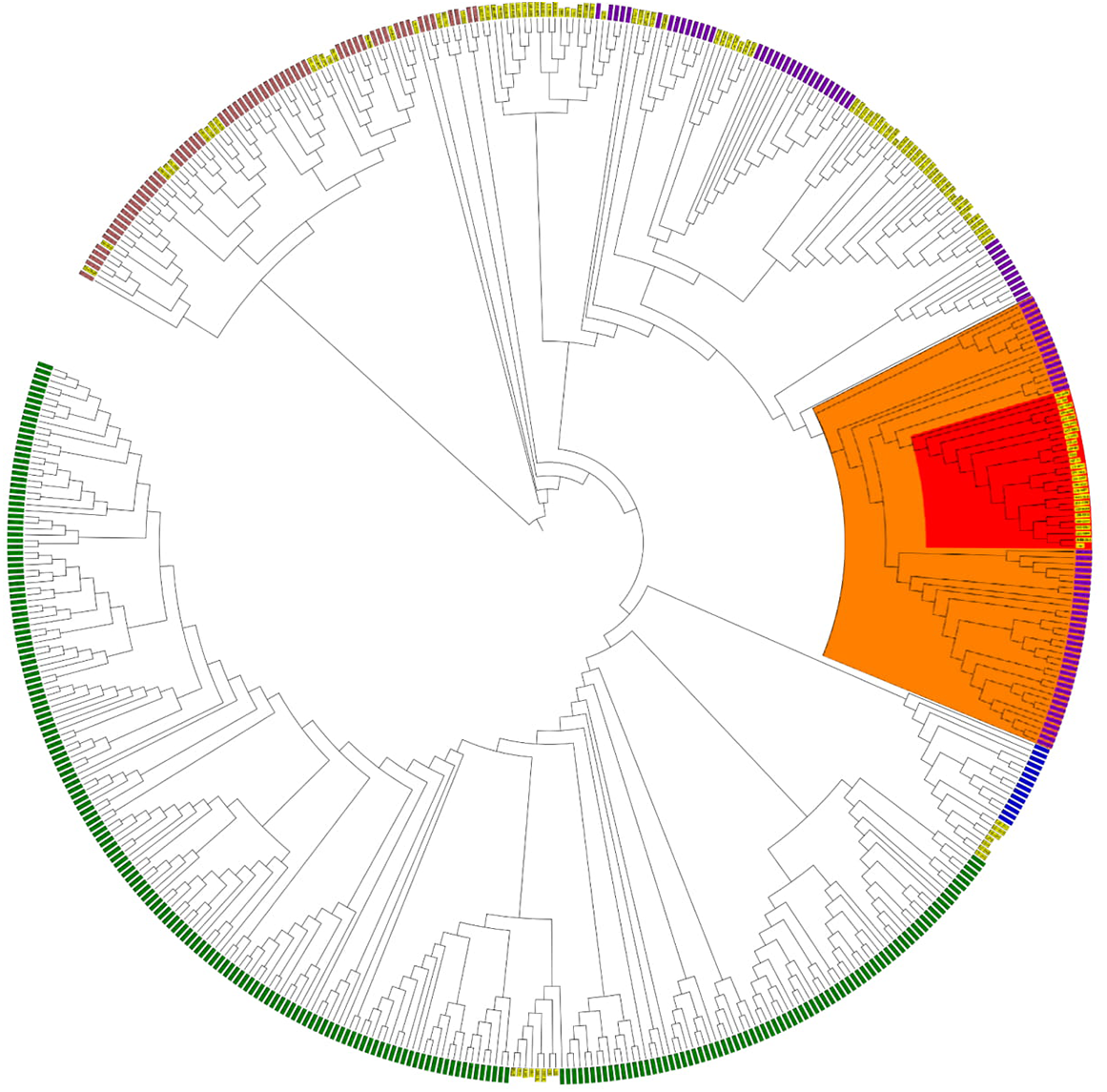
Core genome SNP phylogeny of Australian and international *S. suis* isolates. Pink labels encompass clade 1, purple labels encompass clade 2, blue labels encompass clade 3 and green labels encompass clade 4. Yellow labels indicate Australian isolates. Orange fill encompasses Canadian/Nth American Serotype 2 ST25 and within this red fill encompasses Australian serotype 2 ST25.

**Figure 4.**
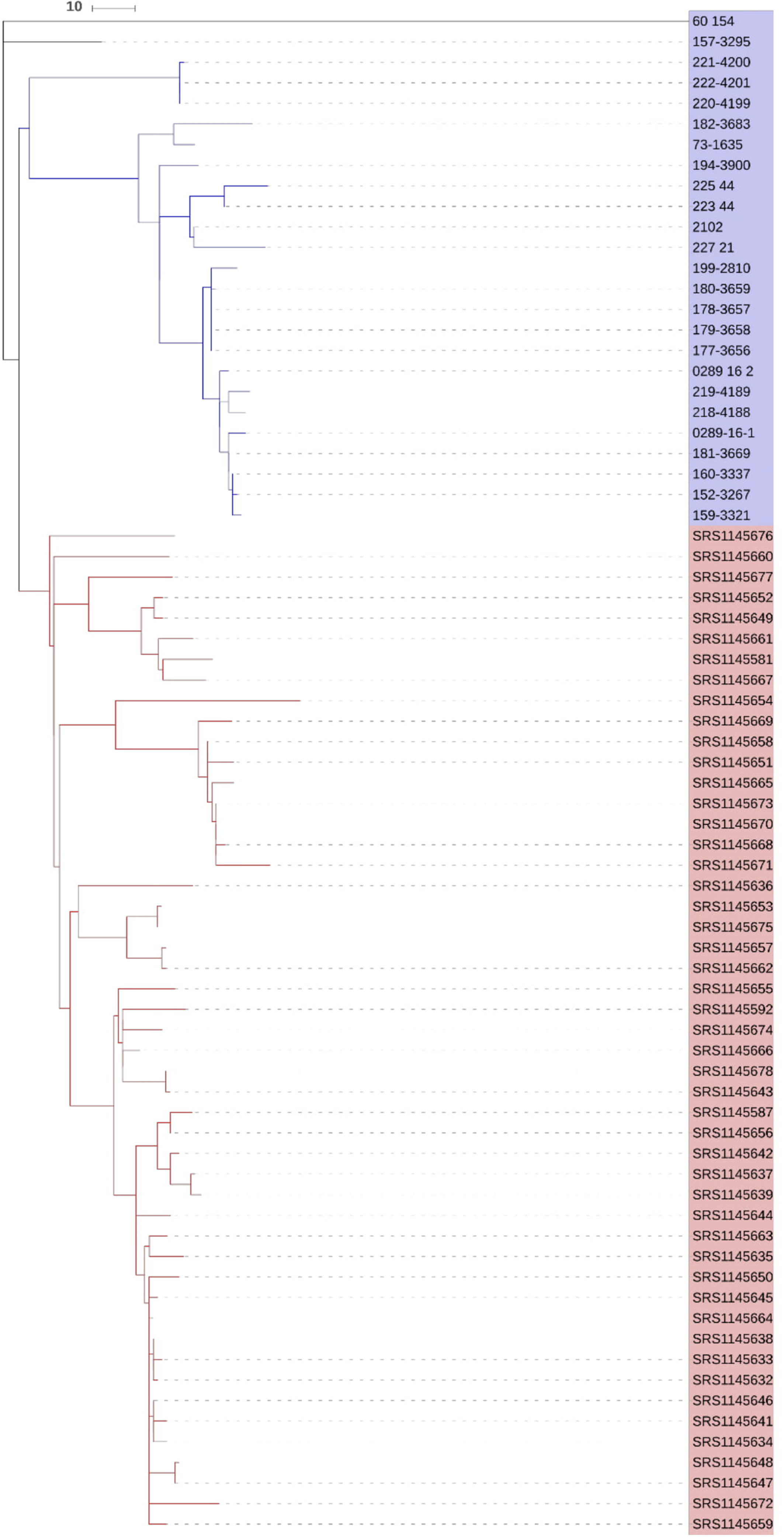
Serotype 2 ST25 isolate core genome SNP phylogeny. Red clade is Canadian/Nth American isolates, blue clade is Australian clade. Remainder are Australian divergent isolates. Scale indicates SNP differences.

### Antimicrobial susceptibility testing

All 148 isolates were subjected to micro-broth dilution to determine the MIC value against ten antibiotics belonging to eight classes (Table 2). A high proportion of the isolates were resistant to both tetracycline (99.3%) and erythromycin (83.8%). Low levels of resistance was observed for florfenicol (14.9%), penicillin G (8.1%), ampicillin (0.7%) and trimethoprim/ sulphamethoxazole (0.7%). None of the isolates were resistant to enrofloxacin.

**Table 2.** Distribution of minimum inhibitory concentrations for *S.* suis. Percentage of isolates classified as non-susceptible (ns ci) and/or clinically resistant (cr ci) with corresponding 95% confidence intervals for those where breakpoints are available. Shaded areas indicate the range of dilutions evaluated. Vertical bars indicate clinical breakpoint (where available). TMS; trimethoprim/sulfamethoxazole

| drug | n | 0.13 | 0.25 | 0.5 | 1 | 2 | 4 | 8 | 16 | 32 | 64 | 128 | 256 | ns_ci | cr_ci |
| --- | --- | --- | --- | --- | --- | --- | --- | --- | --- | --- | --- | --- | --- | --- | --- |
| Ampicillin | 148 |  | 99.3<br>(147) |  |  |  |  |  | .7<br>(1) |  |  |  |  | 0.7(0,3.7) | 0.7(0,3.7) |
| Ceftriaxone | 148 |  | 88.5<br>(131) | 7.4<br>(11) | 2<br>(3) | 1.4<br>(2) |  |  |  |  | .7<br>(1) |  |  | . | . |
| Ciprofloxacin | 148 |  | 37.2<br>(55) | 37.2<br>(55) | 22.3<br>(33) | 3.4<br>(5) |  |  |  |  |  |  |  | . | . |
| Enrofloxacin | 148 | 14.2<br>(21) | 44.6<br>(66) | 40.5<br>(60) | .7<br>(1) |  |  |  |  |  |  |  |  | 0.7(0,3.7) | 0(0,2.5) |
| Erythromycin | 148 | 11.5<br>(17) | 4.1<br>(6) | .7<br>(1) | 1.4<br>(2) |  |  | 2<br>(3) | 1.4<br>(2) | 79.1<br>(117) |  |  |  | 84.5(77.6,89.9) | 83.8(76.8,89.3) |
| Florfenicol | 148 |  |  |  | 2<br>(3) | 4.1<br>(6) | 79.1<br>(117) | 11.5<br>(17) |  | .7<br>(1) | 1.4<br>(2) | .7<br>(1) | .7<br>(1) | 93.9(88.8,97.2) | 14.9(9.6,21.6) |
| Gentamicin | 148 |  |  |  |  | 6.1<br>(9) | 40.5<br>(60) | 43.2<br>(64) | 9.5<br>(14) |  |  |  | .7<br>(1) | . | . |
| PenicillinG | 148 | 81.8<br>(121) | 4.7<br>(7) | 4.7<br>(7) | 3.4<br>(5) | 3.4<br>(5) |  | .7<br>(1) |  | .7<br>(1) |  |  |  | 12.8(7.9,19.3) | 8.1(4.3,13.7) |
| Tetracycline | 148 |  |  |  | .7<br>(1) |  | .7<br>(1) | 1.4<br>(2) | 2<br>(3) | 1.4<br>(2) | 93.9<br>(139) |  |  | 100 (97.5,100) | 99.3(96.3,100) |
| TMS | 148 |  |  | 91.9<br>(136) | 5.4<br>(8) | 2<br>(3) |  | .7<br>(1) |  |  |  |  |  | 8.1(4.3,13.7) | 0.7(0,3.7) |

Among the *S. suis* isolates, 15.6 % were multi-drug resistant (MDR; resistance to ≥3 classes of antimicrobial) (Table 3). Commonly identified MDR phenotypes included macrolide/phenicol/tetracycline (7.4%) and β-lactam/macrolide/tetracycline (6.1%). All isolates carried resistance against at least one antimicrobial class with 73.7% resistant to two classes of antimicrobial classes (2). The most common phenotypic profile was resistance to a combination of macrolides and tetracyclines with 67.6% of isolates determined to carry this resistance. When analysed by clonal type or serotype there was no discernible pattern in AMR phenotype.

**Table 3.** Phenotypic AMR profiles of *Streptococcus suis* isolated from Australian pigs. The breakpoints used to classify isolates in this case are clinical breakpoints. mac; macrolides, tet; tetracyclines, phe; phenicols, bla; beta lactams, fpi; folate pathway inhibitors.

| phenotype | n | % |
| --- | --- | --- |
| 1: mac | 1 | 0.7 |
| 1: tet | 15 | 10.1 |
| 2: mac_tet | 100 | 67.6 |
| 2: phe_tet | 9 | 6.1 |
| 3: bla_mac_tet | 9 | 6.1 |
| 3: mac_phe_tet | 11 | 7.4 |
| 4: bla_fpi_mac_tet | 1 | 0.7 |
| 4: bla_mac_phe_tet | 2 | 1.4 |

The two dominant antimicrobial resistance genes detected in these isolates were *tetO* and *ermB.* The percentage of isolates carrying identified resistance genes for tetracyclines (95.3%) and macrolides (80.5%) support the MIC data with 96.3% and 83.8% of isolates being resistant to tetracyclines and macrolides respectively. Other resistant genes present included aph(3’) (n=6), *fexA* (n=1), *optrA* (n=3), *lnuB* (n=6) and *spc* (n=5).

## Discussion

*Streptococcus suis* is a major cause of disease in pigs, with the potential for significant public health impact. Along with antimicrobial therapy, control measures in swine herds rely on the production of autogenous vaccines, with variable, and often poor, success rates (7, 11). Given the range of serotypes and STs, in conjunction with proven and putative virulence factors and markers, choosing appropriate clones for vaccine stock is difficult, and reliant on having adequate data. In this study we present a detailed assessment of *S.suis* isolates from clinically affected Australian pigs, and compare these isolates to internationally available genome sequence data. Key findings include the virulence potential of ST1 clones, the zoonotic disease potential of Australian ST1 clones which cluster with Vietnamese isolates from cases of human disease, and the limited evolution of Australian clones from their global seed strains.

Analysis of 148 Australian isolates determined that the majority were serotype 2 (26%) or serotype 3 (25%), followed by serotype 1/2 (7.4%). This is consistent with global *S. suis* strains isolated from cases of disease, particularly for serotypes 2 and 3 in Canada, North America and China (8). While the majority of isolates were obtained from lungs, 32 isolates were obtained from brain, abdomen, heart or joint, suggesting invasive or systemic isolates. Of these, 13 were serotype 2 and six were serotype 1/2, however given this relatively small sample size of systemic isolates, an increased pathogenicity cannot necessarily be ascribed to these serotypes. Significantly, even within this sample set, there were 20 individual serotypes detected (Supplementary Table 1). Given that the capsule is a recognised virulence factor and an immunogenic target (12), a minimum requirement for vaccine production should be serotyping from an outbreak strain.

While ST27 and ST25 were the most prevalent MLSTs, 37 individual STs were identified, including 26 new MLSTs. This greatly expands upon previously available Australian data which documented four STs including 1, 25, 28 and 369 (10). The prevalence of isolates as serotype 2 ST25 (17.6%) and serotype 2 ST28 (6.1%) is consistent with those from North America, Canada, Germany, Spain, the United Kingdom and China, and serotype 3 ST27 (18.2%) is consistent with reports from Spain and the United Kingdom (8). These findings indicate that capsular and ST combinations are stable and conserved, and it is likely that the circulating strains present in Australia are reflective of the seed stock which was imported from the United Kingdom, Canada and New Zealand (13).

Despite conservation of serotype/ST combinations, the use of whole genome sequencing demonstrates geographic clustering at the level of the core genome. Analysis of Australian and international isolates demonstrates four clades, although more than 80% of the isolates are present in clades 2 and 4. Clade 2 consists of isolates from the United Kingdom, Canada and Australia. It can be seen that the Australian isolates form distinct clusters, which correlate with serotype/MLST combinations. The most interesting and distinct cluster was seen in Clade 2, whereby Australian serotype 2 ST25 isolates (Fig 4 red) clustered within Canadian serotype 2 ST25 isolates (Fig 4 yellow). Further analysis by re-deriving the core genome for only these isolates showed two distinct populations separated according to geographic origin. This further supports the hypothesis that serotype/ST combinations are stable, however once introduced into a region these will then form divergent populations which over time become phylogenetically distinct from the original clone. Only eight Australian isolates were present in clade 4, which otherwise consisted entirely of Vietnamese and UK isolates. These Australian isolates were all serotype 1/2 ST1: aside from a single isolate which was serotype 14 ST1. This appears to indicate that the MLST of *S. suis* is a better predictor of core genome phylogeny than the serotype. The presence of these isolates and their core genome similarity to the south-east Asian isolates could be due to two reasons. Historically, assays did not differentiate between serotypes 2 and 1/2, with isolates being reported as serotype 2 or serotype 2 (plus 1/2) (14). For this reason it may be that these isolates are derived from European serotype 1/2 isolates which were initially labelled as serotype 2. Another scenario is that a clinically normal swine worker from south-east Asia transmitted a precursor serotype 1/2 clone from the Asian region to Australian pigs. It is difficult to assess the likelihood of this pathway however, given that most human *S. suis* documentation refers to clinically affected humans, clinically affected humans have not been documented to infect other people or pigs, and that the epidemiology and carriage in clinically unaffected humans is not well defined (15).

There are an increasing number of virulence factors termed critical to virulent strains of *S. suis,* which arguably are not critical but in various combinations may affect the virulence of different clones (16). Historically there has been a focus on three particular virulence factors; muraminidase-released protein (*mrp*), extracellular protein factor (*epf*) and suilysin (*sly*), shown to be associated with highly virulent strains, with *cps2* strains carrying all three genes considered the most virulent globally (17). Analysis of Australian isolates revealed that serotype 2 and 3 isolates were almost exclusively *mrp^−^*/*epf^−^*/*sly^−^*. This is in agreement with early studies demonstrating that the presence of all three of these factors is not necessary for a clone to exhibit high levels of virulence (18), with 9/18 (50%) of the isolates obtained from joints or brain having this gene combination. The presence of the *mrp^+^*/*epf^+^*/*sly^+^* combination in 7/11 (63.6%) of serotype 1/2 isolates, along with 4/7 of these isolates being from joints or brain, indicates that Australian serotype 1/2 isolates have particular virulence potential, in contrast to earlier studies which concluded clinical disease in Canadian pigs with serotype 1/2 was likely due to inherent herd factors (19). It was notable that serotype 1/2 isolates were obtained from six separate farms and six separate production enterprises, indicating that this serotype is not confined to a single company or nucleus herd. The mrp^+^/epf^+^/sly^+^ combination was also carried by all ST1 isolates, and these were consistently the isolates with the largest array of putative virulence factors. Like serotype 1/2 isolates, ST1 isolates were obtained from six farms and production enterprises, indicating this is a widespread, potentially virulent clone. The clustering of isolates by MLST is generally reflected in the virulence gene content of isolates as seen in Figure 2, providing further evidence that MLST is a significant indicator of clonal virulence potential when investigating a disease outbreak. In the case of Australian ST1 isolates, a conserved constellation of virulence genes which was not present in other MLSTs provides some insight into the virulence of ST1 clones. While the majority of these genes have their potential virulence characteristics defined in vitro, there is a distinct aggregation of bacterial adhesion factors such as fibronectin binding factors *dpp IV, mrp, epf* and *sbp2* and oligopeptide binding protein *oppA* (20–23). The presence of *ide,* an IgM protease would also be involved in early phase infection and attachment to cells by cleaving IgM blocking cellular binding factors (24).

When this gene block was investigated in ST1 clones from overseas, it was also found to be highly conserved across 226 isolates, with the exception of the *oppA* gene which was absent in 21.2% of isolates. Taken together these factors suggest that both Australian and international ST1 clones have a distinct fitness advantage in terms of binding ability towards host target cells. In order to determine if the potential virulence of Australian ST1 isolates could be classified by analysis of the GDH amino acid sequence, as had been reported in an overseas study, we compared of the amino acid sequence of GDH across the Australian strains (25). We found that all ST1 isolates had amino acid substitutions of A, S, K and K in positions 296, 299, 305 and 330 respectively, the same combination of substitutions reported by Kutz et al. (2008) in highly virulent serotype 2 clones. None of the other Australian STs had this sequence, further demonstrating that virulence may be significantly linked to ST, in this case ST1.

Studies have shown that virulent *S. suis* clones have smaller genome sizes combined with a larger number of virulence genes when compared with less virulent isolates (26). While the sample size of Australian isolates from sites other than the respiratory tract was too small to confirm this, the clustering of ST1 isolates as seen in Figure 2 clearly shows a high number of virulence genes in association with the lowest total gene content of all STs examined. To see if this held true on a global scale, we mapped the same data from 443 isolates which clearly demonstrated the same pattern, inclusive of six ST1 single locus variants. A similar pattern, with clustering in the mid genome size and virulence gene range could be seen with ST25 isolates, and again with a larger total gene number in ST28 isolates. This confers with reports suggesting ST28 clones are non-virulent and ST25 clones are of medium virulence (27) in mouse models. This data provides further evidence that ST1 clones worldwide carry high virulence gene content in combination with a low total number of genes and that virulence is more closely associated with MLST than cps type (28).

Antimicrobial resistance of Australian strains was similar to levels reported overseas with regards to tetracycline (99.3%), erythromycin (83.8 %) and trimethoprim/sulfamethoxazole (0.7%). Resistance to florfenicol was 14.9%, while all isolates were clinically susceptible to enrofloxacin, likely due to this being banned from use in food producing animals in Australia, and evidence of the success of this programme. Of concern was the observed clinical resistance to penicillin G, albeit at a relatively low level in 8.1% of isolates. This is a first line therapy for *S. suis,* and indeed the beta-lactams are used in human therapy (29). Penicillin resistance levels of 5% have been reported in isolates from England (30), and 0% in isolates from China (31). Therefore this is an aspect of *S suis* in Australia that must be carefully monitored from both an animal and public health point of view. It should also be noted that our analysis did not detect any common beta-lactamase genes, as was the case in a recent study of Spanish and Canadian serotype 9 isolates (32), indicating a potentially unknown mechanism of resistance in these isolates.

This study has greatly increased the data available on the circulating strains of *S. suis* in the Australian pig herd. Comparative genome analysing using Australian and overseas data revealed that production of autogenous vaccine stock should not be based totally on serotype of circulating strains. In fact it is our view that it is equally important to take into account MLST, as we have demonstrated that there is a strong correlation between MLST and virulence gene content. It is possible that the core genome, or specific gene constellations associated with MLST groupings as seen with ST1 isolates in this study, may uncover suitable vaccine targets for development of pan-MLST vaccines. Therefore, we recommend that the MLST of isolates from clinical cases be regularly monitored to ensure they match vaccine stocks. With the rapidly expanding range of whole genome sequence analysis pipelines such as that used in this study, clonal typing and virulence marker determination of clones associated with disease outbreaks can be achieved within a matter of days. Utilisation of this technology can allow for highly targeted autogenous vaccine use. Additionally, the presence of bacterial adhesion factors associated with ST1 clones which were not present in other clonal groups also presents a target for future vaccine studies, potentially in the form of multivalent subunit vaccines.

Additionally, this study demonstrates the phylogenetic relationship of *S. suis* clones evolving from single clonal types imported into a country. Australia’s animal import policies have prevented live pigs from being imported since 1987 (13), meaning that circulating clones are a derivation of those from at least 30 years ago and likely introduced with stock from the United Kingdom, Canada and New Zealand. As can be seen in clade 1 of Figure 3, approximately 13% (20/148) of Australian clones, only one of which was from brain tissue, have diverged significantly from the majority of those studied, and consist predominantly of serotypes 16 and 31, with all except isolate 74-1662 having newly assigned MLSTs. Despite this, it can be seen that for the majority of isolates the core genome has remained consistent with European and North American isolates, such that while Australian clones evolved in isolation, they have continued clustering closely with their hypothesised genetic predecessors.

In conclusion, Australian clones associated with disease in pigs consist predominantly of serotypes 2, 3 and 1/2, which is consistent with reports from other pig producing countries (8). Despite the limited number, the characterisation of serotype 1/2 ST1 clones is significant, as all displayed distinctive factors associated with highly virulent *S. suis,* along with grouping separately to other Australian isolates in the Vietnamese/UK clade, potentially indicating a separate source of introduction. In addition to this, these strains have only low levels of divergence from Vietnamese and UK isolates from cases of swine and human systemic disease, making this a sequence type which requires further investigation.

## Methods

### Isolates

Swabs or fresh tissue from diseased pigs were submitted by consultant veterinarians to ACE Laboratory Service, Victoria, Australia. Samples were plated onto Sheep Blood Agar (SBA), MacConkey Agar and Chocolate Agar (Oxoid, Thermo Fisher Scientific), and incubated at 37°C for 24 hours. Suspect *S. suis* isolates, which showed alpha haemolysis on SBA and that were assessed to demonstrate morphology consistent with *S. suis* were sub-cultured onto fresh plates for isolation. Identification was carried out using matrix assisted laser desorption ionization-time of flight mass spectrometry (MALDI-TOF) typing (Bruker). Following confirmation, isolates were harvested from SBA plates into 1ml of Tryptic Soy Broth (TSB)+10 % glycerol and stored at −80°C in sterile microcentrifuge tubes. A random subset of isolates was thawed and streaked on to SBA for resurrection at 37°C. Purity of each isolate was confirmed prior to harvesting plates onto transport swabs for postage to Murdoch University.

### MIC Testing

All isolates were subjected to antimicrobial susceptibility testing via broth microdilution according to the Clinical Laboratory Standards Institute (CLSI) Performance Standards. MIC results were categorized as susceptible, intermediate and resistant using the clinical interpretative criteria specified in CLSI performance standard VET01-S3 (33). If interpretive criteria was not present in VET01-S3, CLSI performance standard M100-S25 was used (34).

### Whole-genome sequencing

DNA extractions were performed on the 148 isolates using a MagMax DNA multi-sample kit (ThermoFisher Scientific) according to the manufacturer’s instructions, with the modification to omit the RNAse treatment step. Library preparation was performed with a Nextera XT kit with the only change from the manufacturer’s instructions being an increased tagmentation time of 7 minutes. Sequencing was performed on an Illumina Nextseq 500 platform using a mid-output V2 (2 x 150 cycles) reagent kit.

### Sequence analysis

All sequencing files were parsed through the Nullarbor bioinformatics pipeline (v1.20) (35) to determine MLST and antimicrobial resistance genes. A database was manually created containing all capsular serotypes and 40 previously described putative virulence genes (36). The Abricate programme was used to query contig files against the database to determine capsular type and virulence genes, with cutoffs of ≥95% coverage and ≥99% identity used to determine gene presence. Distinguishing serotype 2 from serotype 1/2, and serotype 1 from serotype 14 was performed as outlined by Athey et al. (2016) (37).

Phylogenetic trees were based on single nucleotide polymorphisms in the core genome. For genomic comparison against international *S. suis* isolates, 383 previously published sequences from Vietnamese, UK, North American and Canadian isolates (26, 38)were downloaded from NCBI and ENA. Genome annotation was performed using Prokka (v1.12) (39) and outputs were processed using Roary (v3.8.0) (40) for core genome determination and Gubbins (v2.2.3) (41) for recombination removal and alignment. Manual annotation of trees was performed in iTOL (v4.2) (42). Sequences for new MLST allele variations were uploaded to the *Streptococcus suis* MLST Databases site (https://pubmlst.org/ssuis/) for assignment of allele identification, and final allele combinations were then uploaded for assignment of new MLSTs.

## Accession number(s)

All sequence read data generated in this study has been deposited in the NCBI Sequence Read Archive under accession number SRP150885.

---

**Supplementary Table 1**. Summary table of isolate metadata from study.

